# Accurate alignment between correlated light, cryo-light, and electron cryo-microscopy data using sample support features

**DOI:** 10.1101/099077

**Authors:** Karen L. Anderson, Christopher Page, Mark F. Swift, Dorit Hanein, Niels Volkmann

## Abstract

Combining fluorescence microscopy with electron cryo-tomography allows, in principle, spatial localization of tagged macromolecular assemblies and structural features within the cellular environment. To allow precise localization and scale integration between the two disparate imaging modalities, accurate alignment procedures are needed. Here, we describe a marker-free method for aligning images from light or cryo-light fluorescence microscopy and from electron cryo-microscopy that takes advantage of sample support features, namely the holes in the carbon film. We find that the accuracy of this method, as judged by prediction errors of the hole center coordinates, is better than 100 nm.

## Introduction

Correlative imaging of samples using fluorescence markers to identify macromolecular assemblies and structural features in images and volumes obtained by electron cryo-microscopy approaches has become an important tool for biomedical research in recent years. In electron cryo-microscopy, samples are vitrified by fast freezing, resulting in the biological samples being embedded in a layer of amorphous ice, preserved in their native state (Dubochet et al., 1988). Recent hardware and software developments have revolutionized the field, allowing now to obtain structural information at near atomic resolution of a wide variety of macromolecular assemblies (Subramaniam et al., 2016) and also enabling imaging of unperturbed cellular landscapes at high resolution (Asano et al., 2015). Although some large macromolecular assemblies are identifiable in such cellular reconstructions (Asano et al., 2015; Mahamid et al., 2016; Woodward et al., 2015) the majority of the components are not identifiably without specific labeling, presenting a major challenge in crowded cellular environments.

The advantages of fluorescence microscopy are complementary to those of electron cryo-microscopy. The fluorescence microscopy approach allows live cell imaging, following specific fluorescently labeled molecules in real time, and the larger field of view facilitates the identification and location of sparse cellular events in a larger context. The combination of the two microscopy approaches potentially allows identifying regions associated with specific biological events by fluorescence microscopy and then investigating the underlying structural features at high resolution by electron cryo-microscopy.

For correlative imaging, vitrification of the sample can be either performed before or after fluorescence imaging. The advantages of room temperature fluorescence imaging include the use of optimized equipment such as high-powered, oil-immersion lenses with large numerical apertures and that dynamic processes can be characterized and followed. Even if precautions are being made to avoid imaging induced damage caused by buffer and anti oxidative agents, the downside of this procedure is that structures may change in the time required to perform the vitrification. To avoid this problem, cells can be fixed. For carefully chosen fixation protocols, although they might introduce minor artifacts through cross-linking of cellular components, most structural relationships and components are well preserved (Briegel et al., 2010). For vitrification before fluorescence imaging, specialized cryo-stages that allow fluorescence imaging at cryo-temperatures are required (Sartori et al., 2007; Briegel et al., 2010; Faas et al., 2013; Hampton et al., 2017) and the resolution is restricted to about 400 nm or less owing to lack of immersion cryo-objectives. In addition, cryo-fluorescence imaging loses the capability of capturing the dynamics of the biological process. Correlative imaging using super-resolution techniques and electron cryo-microscopy are also under development (Wolff et al., 2016).

For all these correlative techniques, it is necessary to find an accurate alignment between the images from the two imaging modalities. The required coordinate transform generally involves rotation, translation, and scaling. In some instances, when there are significant beam-induced changes to the sample (Brilot et al., 2012) or the sample support is tilted in respect to the second imaging modality, further correction may be necessary. In practice, the latter appear to be of minor consequence (Schellenberger et al., 2014). Early applications of correlative fluorescence electron cryo-microscopy imaging took advantage of large cellular features, which allowed a straightforward relative orientation of the sample (van Driel et al., 2009). To increase the accuracy of the alignment and to allow determination of accuracy estimates, the use of fluorescence beads as fiducial markers for alignment was introduced (Schorb and Briggs, 2014; Schellenberger et al., 2014). Here, we propose a marker-free approach using features of the sample support to provide alignment between images from fluorescence or cryo-fluorescence microscopy and electron cryo-microscopy images. The alignment quality of this approach is with 50-100 nm estimated accuracy comparable to that previously reported for marker-based alignment and does not involve modification of the experimental procedures.

## Materials and methods

### Sample preparation

GFP-transfected mouse embryonic fibroblast and Chinese hamster ovary cells (Bachir et al., 2014) were cultured on electron microscopy grids as previously described (Anderson et al., 2016). *Streptococcus pneumoniae* samples were prepared as previously described (Durand et al., 2015). Grids were manually plunge frozen in liquid ethane at liquid nitrogen temperature either after room temperature fluorescence imaging or before cryo-fluorescence imaging.

### Correlative fluorescence and electron cryo-microscopy

We tested the alignment protocols using samples on commonly used electron microscopy grids (specialty grids 300, 400, and 200 mesh copper, Quantifoil Grids or C-Flat with 1.2/1.3, 2/1, 3.5/1, 2/2 hole patterns). Fluorescence modalities used for correlated imaging included FRET, TIRF, confocal, and cryo-fluorescence imaging with mCherry or GFP. Cryo-fluorescence imaging was performed using a CorrSight CLEM microscope (FEI Company), equipped with a motorized stage, a broad-band Xenon light source and motorized fluorescent filter sets. The vitrified grids were mounted into grid support rings (Auto-gridTEM sample holder, FEI) and transferred into the cartridge holder of a Titan Krios (FEI company). Electron cryo-microscopy images were taken with an FEI Titan Krios transmission electron cryo-microscope equipped with an extra-high brightness field emission gun (XFEG) and operated at 300 kV using a back-thinned 4k × 4k FEI Falcon II direct detection device.

### Identification of holes in the carbon film

To obtain estimates for the center positions of the holes in the carbon images in the electron cryo-microscope, we first applied an iterative median filter (van der Heide et al., 2007) to suppress background noise. We then applied the Canny edge detector (Canny, 1986), and run a circular Hough transform (Kimme et al., 1975) with variable radius to determine the optimal radius for detecting the holes. The center positions of the detected circles detected with the optimal Hough-transform radius were stores as references for aligning the fluorescence images. To extract the circle centers from the fluorescence images, we increased the contrast to maximum and run an iterative median filter to suppress the noise. The resulting images were subjected to Canny edge detection and circular Hough transform as described above for the electron cryo-microscopy images. The center coordinates of the detected circles were extracted for further processing.

### Alignment using the circle center coordinates

To allow the use of the circle center coordinates for alignment, we first determined the mapping of the same circles in the two different imaging domains. After manual assignment of the correspondences, a least-squares coordinate fitting approach (Kabsch, 1976) was used to determine the relative rotation, translation and accurate scales between the images.

### Alignment accuracy estimation

To obtain an estimate of the alignment accuracy, we averaged over individual prediction errors following a similar procedure previously developed for fiducial-based alignments (Schellenberger et al., 2014). For the calculation of individual prediction errors, one hole center coordinate from the fluorescence image was excluded from the alignment calculation. This coordinate was treated as the object of interest and its position predicted within the electron cryo-microscopy image. The predicted position was compared to the actual position in the electron cryo-microscopy image to estimate the individual prediction error. This operation was repeated for all hole centers in the fluorescence images. The estimate of the alignment accuracy was then calculated as the average of the individual prediction errors.

## Results

We developed a method for accurate alignment of fluorescence images from light microscopy with images obtained by electron cryo-microscopy. The approach is based on the visibility of holes in the carbon support in both imaging modalities. The holes are readily identifiable in the electron cryo-microscopy images (Figure 1A) but in the fluorescence images the holes are not immediately apparent (Figure 1B). However, there is always a subtle difference between the background fluorescence in the holes versus fluorescence on the carbon that becomes visible when increasing the contrast to maximum. The resulting images tend to be rather noisy, which makes accurate detection of the hole boundaries difficult. Application of several rounds (3-8) of iterative median filtering (van der Heide et al., 2007) improves the situation considerably and most holes become identifiable in the fluorescence images as well (Figure 1C). The electron cryo-microscopy images also benefit from a few rounds of iterative median filtering, which suppresses the background variations, especially inside the holes. A combination of edge detection (Canny, 1986) and circular Hough transform (Kimme et al., 1975) to the enhanced images allows accurate determination of the hole sizes and boundaries, and thus their center coordinates, in both imaging modalities (Figure 2A-B, red circles). It is noteworthy that the circular Hough transform is well suited to complete circles that are partially obscured, allowing to remove possible bias by obstructions that behave different in the two imaging modalities.

**Figure 1:**
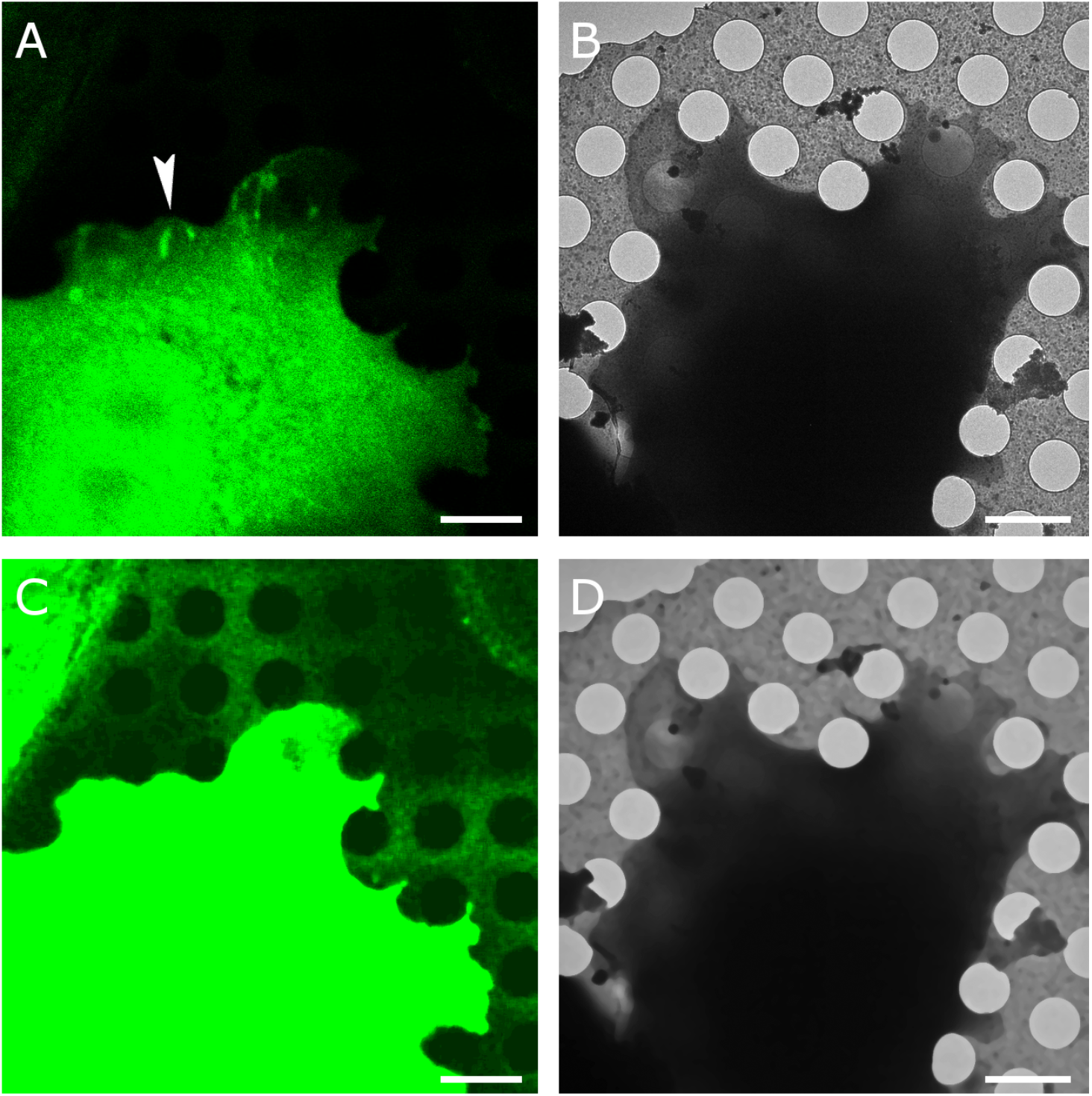
Correlative imaging. **A**. Correlated Paxillin-GFP TIRF fluorescence (green) and **B**. electron cryo-microscopy imaging of the same cell. The arrowhead in A points at a region with elongated spots of high paxillin fluorescence, indicative of focal adhesions, an actin-based assembly involved in cell adhesion to extracellular substrates. **C**. The holes in the carbon film are not visible in A but become visible after contrast adjustment and noise suppression with iterative median filtering. **D**. Iterative median filtering also suppresses background fluctuations in the electron cryo-microscopy image, allowing more accurate determination of the hole centers. Bars are 2 microns.

**Figure 2:**
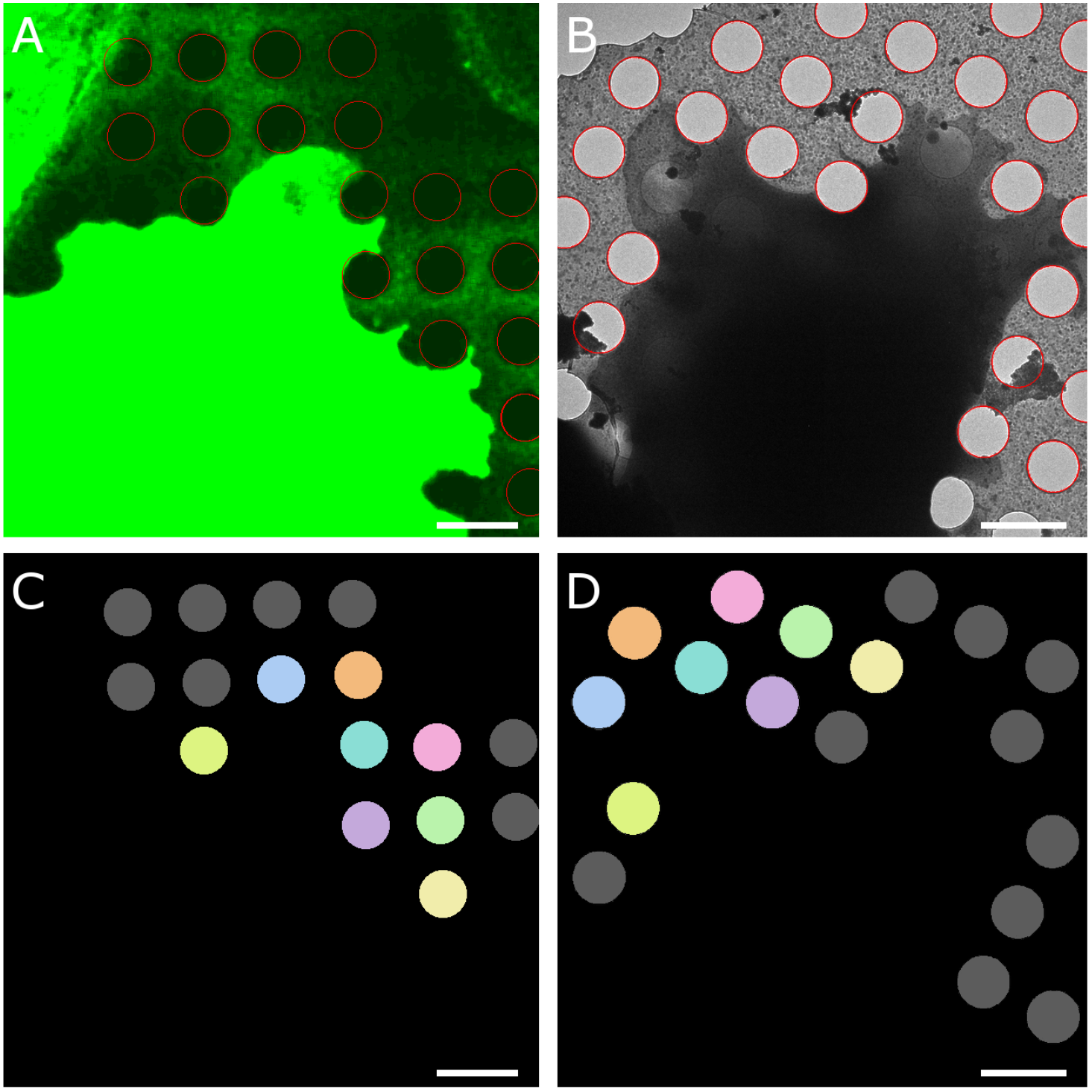
Hole detection. **A**. Application of edge detection and circular Hough transform allows localization of the hole boundaries (red circles) in the enhanced fluorescence image. **B**. Hole boundaries (red) in electron cryo-microscopy image of the same cell. **C-D**. Correspondence mapping of the holes in the respective images. Identical colors refer to the same hole, holes marked in grey do not have a corresponding hole in the other image modality. Bars are 2 microns.

Once the hole center locations are determined, a one-to-one correspondence mapping between the holes in one versus the other imaging modality is determined (Figure 2C-D). The coordinates can then be used to determine the rotation, translation, and relative scale between the fluorescence and electron cryo-microscopy images. A least-square approach (Kabsch, 1976) is accurate and fast. To estimate the alignment accuracy, we calculated averages of the prediction errors of individual circle centers {Schellenberger 2014). We find that the average root-mean-square deviation between the predicted coordinates and the coordinates based on the full set is between 50 and 100 nm, depending on the actual correlative image pair. The standard deviations of the alignment accuracy estimates were generally below 10% of the estimate. Application of the coordinate transform to the fluorescence images shows excellent visual overlap between the hole boundaries with the electron cryo-microscopy images (Figure 3A) and allows accurate overlay of features of interest in the fluorescence image with the electron cryo-microscopy image of the same region (Figures 3B, 4).

**Figure 3:**
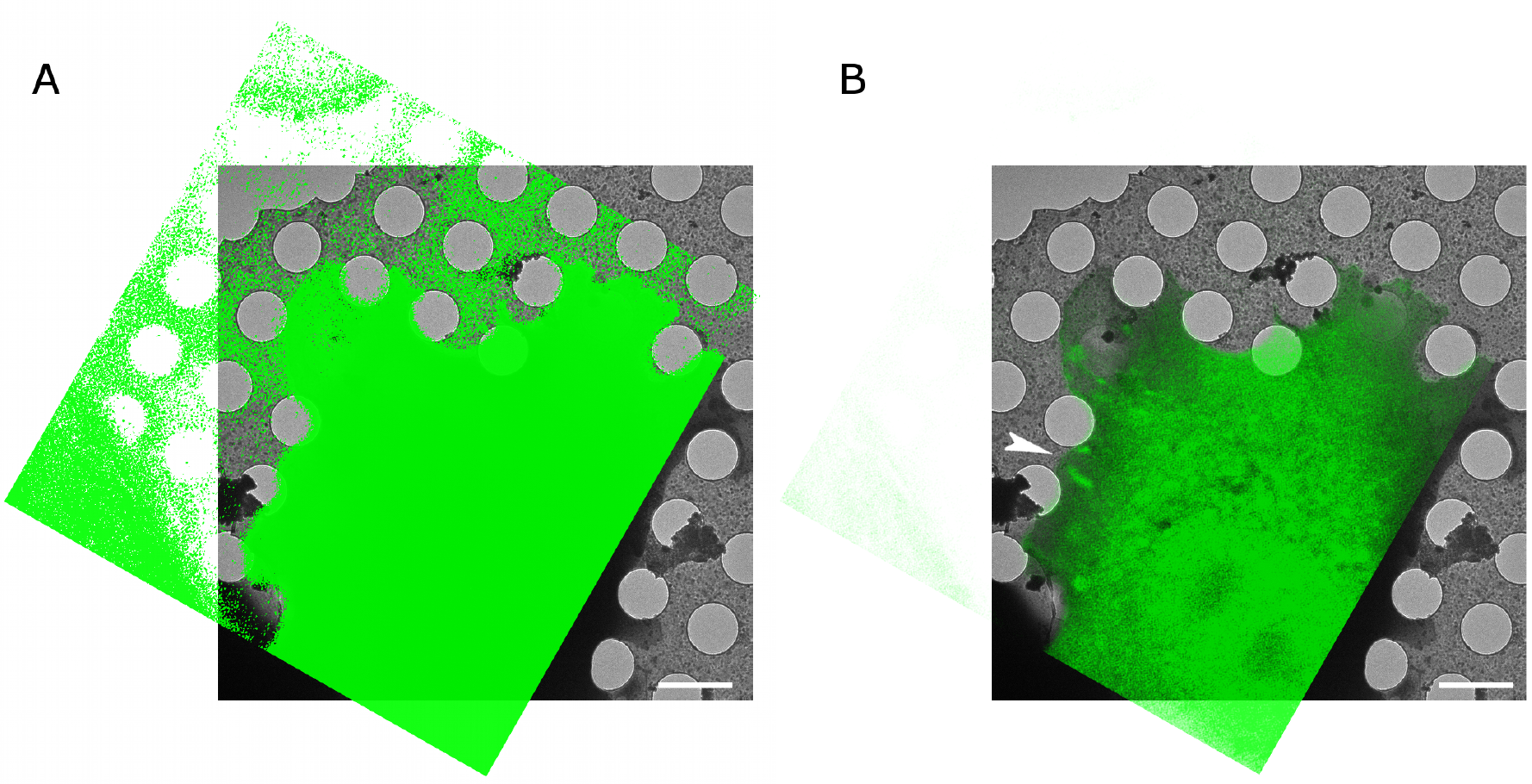
Overlay of Paxilin-GFP fluorescence image (green) with electron cryomicroscopy image (greyscale) after applying the transform determined using the carbon hole center correspondences. **A**. Is shown at high contrast to show the excellent correspondence between the holes. **B**. Is shown at lower contrast to show the overlay of fluorescence features with the electron cryo-microscopy image. The arrowhead points at the same focal adhesion region as the arrowhead in Figure 1. The bars are 2 microns.

**Figure 4:**
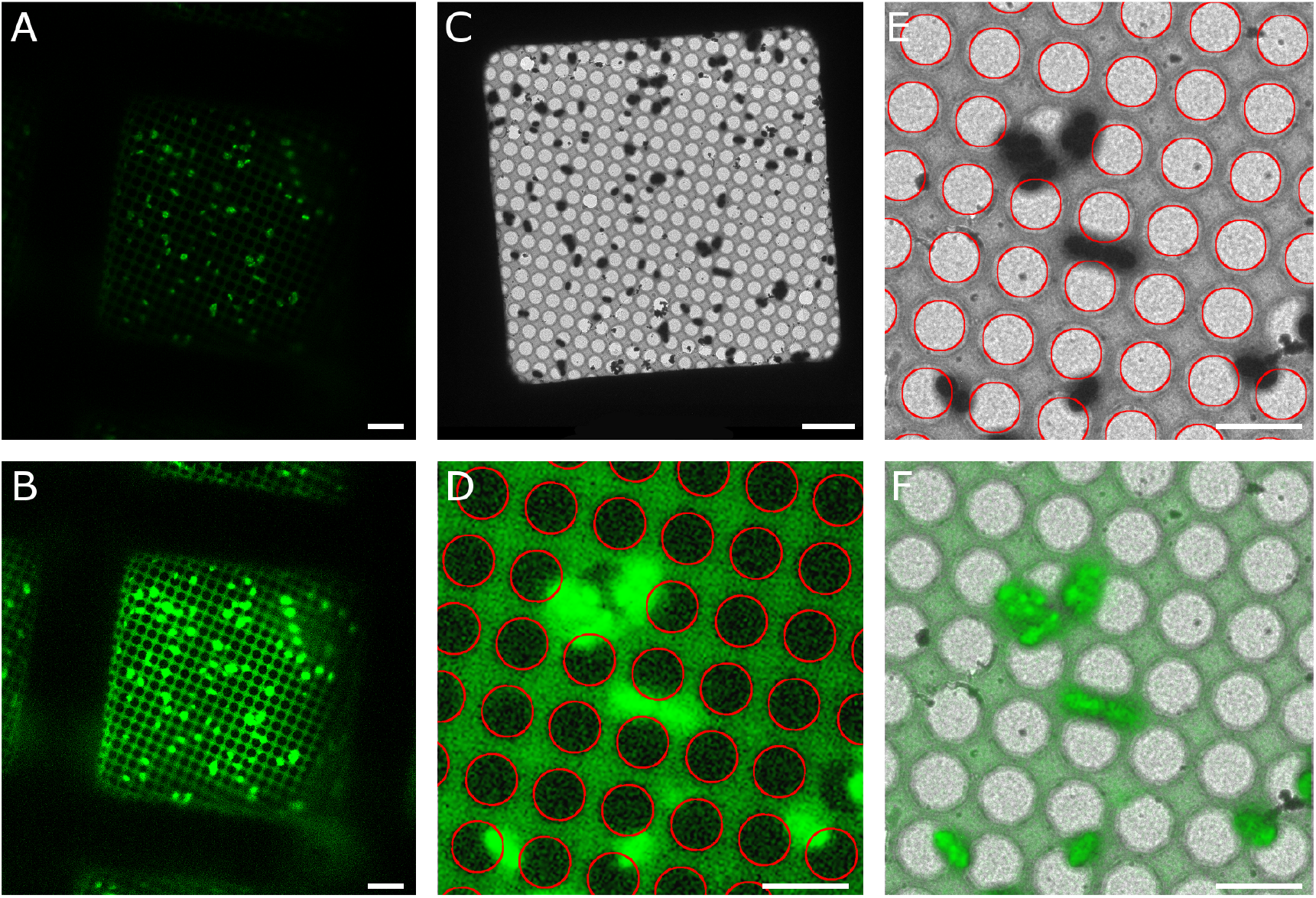
Correlating cryo-fluorescence and electron cryo-microscopy images. **A**. Correlated R3045 ComGA:GFP fluorescence (green) of expressing *Streptococcus pneumoniae* recorded with a CorrSight cryo-microscope (FEI company). **B**. Enhanced cryo-fluorescence image showing the detectability of the holes in the carbon film. **C**. Electron cryo-microscopy image of the same field of view. **D.** Enlarged region of enhanced cryo-fluorescence with Hough transform circles (red) overlaid. **E**. Enlarged region of electron cryo-microscopy with Hough circles (red) overlaid. **F**. Alignment of cryo-fluorescence (green) with electron cryo-microscopy image (grayscale) based on the centers of corresponding Hough circles. Bars are 5 microns.

## Discussion

Fiducial-less alignment based on features of the electron-microscopy sample support has several advantages over the more traditional marker-based alignment methods. First, no additional material needs to be added to the sample. Markers need to be explicitly introduced and may obscure features of interest. In contrast, the sample support is always present. Second, the distribution of the holes is regular and covers the entire grid in predictable ways with excellent coverage. Marker concentrations need to be optimized. Too many markers can interfere with the analysis of the sample; too few markers may not be enough for high-quality alignment. In addition, markers can spatially cluster, which can also degrade alignment quality. For example, if ten markers form a small cluster in the lower right-hand corner, then alignment between features in the top left-hand corner may be off significantly even though the precision estimate using the ten marker coordinates will not give any indication of the issue. Third, the large size of the holes makes them amenable for aligning over a larger range of magnifications than point-like fiducial markers such as fluorescent beads can provide.

The estimated alignment accuracy of our method is with 50-100 nm comparable to accuracy estimates previously provided for marker-based alignment of fluorescence and electron cryo-microscopy images (Schellenberger et al., 2014; Schorb and Briggs, 2014). For bead-based alignment, it is assumed that the bead is essentially a point-like object at the light-microscopy level and a two-dimensional Gaussian is fitted to get a precise, sub-pixel estimate of the center position of the bead (Schellenberger et al., 2014). When using holes for alignment, a circle must be fitted to the rim of the hole to determine the precise center position. On one hand, this can be an advantage because more image pixels are available for fitting the circle than are normally available for fitting a two-dimensional Gaussian to a bead position. On the other hand, because of the relatively large size of the circles, small errors in center estimates might not be as apparent as an offset Gaussian would be. The cumulative effect does not appear to make a significant difference because the overall alignment accuracy of bead-based and hole-based alignments is comparable.

The use of the holes in the carbon film of the electron cryo-microscopy sample support for alignment is a viable alternative to bead-based alignment techniques in terms of alignment accuracy and does not require any modification of the sample and is thus less invasive than bead-based methods. The regular and predictable distribution of holes ensures adequate coverage of reference points for alignment in any given area on the grid and the regularity will also help to detect and correct for distortions and relative tilt of the sample support between imaging modalities.

## Acknowledgements

This work was supported by NIGMS grant R01 GM115972 and CCSG Pilot Project funding (P30 CA030199). We thank Drs. Rémi Fronzes and Esther Marza for providing the R3045 ComGA:GFP *Streptococcus pneumoniae* samples.

